# Culexarchaeia, a novel archaeal class of anaerobic generalists inhabiting geothermal environments

**DOI:** 10.1101/2022.04.06.487207

**Authors:** Anthony J. Kohtz, Zackary J. Jay, Mackenzie Lynes, Viola Krukenberg, Roland Hatzenpichler

## Abstract

Geothermal environments, including terrestrial hot springs and deep-sea hydrothermal sediments, often contain many poorly understood lineages of archaea. Here, we recovered ten metagenome-assembled genomes (MAGs) from geothermal sediments and propose that they constitute a new archaeal class within the TACK superphylum, “*Candidatus* Culexarchaeia”, named after the Culex Basin in Yellowstone National Park. Culexarchaeia harbor distinct sets of proteins involved in key cellular processes that are either phylogenetically divergent or are absent from other closely related TACK lineages, with a particular divergence in cell division and cytoskeletal proteins. Metabolic reconstruction revealed that Culexarchaeia have the capacity to metabolize a wide variety of organic and inorganic substrates. Notably, Culexarchaeia encode a unique modular, membrane associated, and energy conserving [NiFe]-hydrogenase complex that potentially interacts with heterodisulfide reductase (Hdr) subunits. Comparison of this [NiFe]-hydrogenase complex with similar complexes from other archaea suggests that interactions between membrane associated [NiFe]-hydrogenases and Hdr may be more widespread than previously appreciated in both methanogenic and non-methanogenic lifestyles. The analysis of Culexarchaeia further expands our understanding of the phylogenetic and functional diversity of lineages within the TACK superphylum and the ecology, physiology, and evolution of these organisms in extreme environments.

## Introduction

In the past two decades, massive efforts in recovering metagenome assembled genomes (MAGs) from environmental samples have resulted in the description of many novel high-ranking archaeal lineages (1, 2). The current picture of archaeal diversity comprises four superphyla - the Asgard, DPANN, *Euryarchaeota*, and TACK archaea - and it is likely that high-ranking lineages (phylum, class, order) are yet to be discovered (1, 2). Many archaeal phyla, including *Nanoarchaeota* (3), *Ca.* Korarchaeota (4), *Ca.* Geoarchaeota (5), *Ca.* Odinarchaeota (6), *Ca.* Marsarchaeota (7), *Ca*. Nezhaarchaeota (8), and *Ca.* Brockarchaeota (9), were originally discovered in extreme geothermal habitats. While most of these lineages have no cultured representatives, they are often proposed to play important roles in biogeochemical cycles (2, 10).

In lieu of cultures, many lineages are currently best understood via MAGs, which allow for determining their taxonomic placement, inferring their metabolic potential, and generating hypotheses on their ecophysiology that may lead to subsequent *in situ* studies or eventual cultivation (1, 11). For example, metagenomics revealed that several lineages within the TACK superphylum encode methyl-coenzyme M reductase (MCR), a hallmark enzyme long-thought to be exclusive to alkane-cycling *Euryarchaeota*. This suggests that the potential for methanogenesis and alkane oxidation are more widespread within the archaeal domain than previously realized (8, 12–15). Additionally, the potential for anaerobic methylotrophy, a process that in archaea had been solely described in methylotrophic methanogens, was recently detected in MAGs of non-methanogenic *Ca.* Brockarchaeota (9). Considering the large diversity of uncultivated taxa revealed by environmental metagenomics, characterizing new archaeal lineages through MAGs is crucial to understanding the phylogenetic and metabolic diversity that exists among archaea and revealing their potential impacts on biogeochemical cycling and ecosystem functioning (16).

Here, using ten MAGs from terrestrial hot spring and deep-sea hydrothermal seep sediments, we report on a new archaeal class, “*Candidatus* Culexarchaeia”. We describe their biogeographic distribution across geothermal systems, evaluate their phylogenetic affiliation through marker gene and functional gene analysis, and describe genes encoding unique protein complexes and versatile metabolic pathways.

## Materials and methods

### Sample collection, DNA extraction, and metagenome sequencing

Two hot springs located in the Lower Culex Basin (LCB) thermal complex of Yellowstone National Park (YNP) were sampled for molecular and geochemical analyses. At the time of sampling in October 2017, YNP site LCB-003 (44.57763, −110.78957) had a temperature of 72.5 °C and a pH of 6.47, while YNP site LCB-024 (44.57347, −110.79504) had a temperature of 69.4 °C and a pH of 7.79. Surface sediments (∼1 cm deep) were collected, immediately frozen on dry ice before transfer to the lab, and subsequently stored at −80°C until further processing. DNA was extracted from ∼1 g of sediment using the FastDNA Spin Kit for Soil (MP Biomedicals) according to manufacturer’s instructions (MP Bio). DNA extracts were shotgun sequenced at the Joint Genome Institute (JGI). Truseq libraries were prepared using low input (10 ng for LCB-003) or regular input (100 ng for LCB-024) quantities of DNA. Libraries were sequenced on an Illumina NovaSeq platform using the NovaSeq XP V1 reagent kits, S4 flow cell, following a 2×150 indexed run recipe.

### Metagenome assembly, binning, and quality assessment

Metagenomic reads for YNP sites LCB-003 and LCB-024 metagenomes were quality filtered according to JGI’s analysis pipeline and assembled with SPAdes v3.12.0 (17) with the following settings: -k 33,55,77,99,111 -meta. Assemblies for the Great Boiling Spring (GBS) (18), Washburn Hot Springs (WB) (14), and Guaymas Basin (GB) samples were downloaded from the IMG/M portal (19). Assembled scaffolds ≥ 2,000 bp for each metagenome were binned using six different approaches with four different programs, Maxbin v2.2.4 (20), Concoct v1.0.0 (21), Metabat v2.12.1 (with and without coverage) (22), and Autometa v1 (bacterial and archaeal modes, including the machine learning step) (23). Bins produced from each program were refined with DAS_Tool (24). In addition, the published MAG JZ-Bin-30, recovered from a metagenome obtained from a geothermal well in the Yunnan province (China) (25, 26), was downloaded from the IMG/M portal. CheckM (27) was used to estimate MAG completeness, redundancy, and relative abundance. tRNAs were identified with tRNAscan-SE (28), using the archaeal-specific covariance models. Optimal growth temperature was predicted for each MAG using Tome, which analyzes proteome-wide 2-mer amino acid compositions (29).

### Geochemical analyses

Geochemical data for most sites discussed in this study have been described previously: GBS (18), WB (14), and Jinze Spring (JZ) (25, 26). The Guaymas Basin (GB) metagenomes were limited in available geochemical data, but the temperature range was 53-83 °C near the time of sampling, based on the temperature profiles measured near the position the sediment core was collected (Supplementary Table 1). For YNP sites LCB-003 and LCB-024 samples for aqueous geochemistry, including dissolved oxygen, dissolved sulfide, dissolved iron, and dissolved gasses (H2, CH4, and CO2) were collected as described previously (30) and are available in Supplementary File 7.

### Phylogenetic analyses

The 16S rRNA gene sequences encoded in Culexarchaeia MAGs were used in BLASTn searches to screen NCBI and IMG databases for related sequences. Culexarchaeia 16S rRNA sequences were aligned against reference archaeal 16S rRNA sequences and masked using SSU-ALIGN (31), which produced a final alignment of 1,376 positions that was used for phylogenetic analyses. Maximum likelihood analysis was performed using IQtree2 (32) v2.0.6 with the nucleic acid model GTR+F+G and 1,000 ultrafast bootstraps. A set of 43 single-copy marker proteins (Supplementary File 1) used in a previous phylogenomic study (7), were collected from Culexarchaeia MAGs and archaeal reference genomes. These markers were aligned with MUSCLE (33), trimmed with trimAL (34) using a 50% gap threshold, and concatenated. The final trimmed and concatenated alignment of 10,578 positions was used for phylogenetic analysis. Maximum likelihood analysis was performed using IQ-tree2 (32) v2.0.6 with the LG+C60+F+G model and 1,000 ultrafast bootstraps. Additionally, a set of 46 ribosomal proteins (listed in Supplementary File 1) found in Culexarchaeia MAGs and archaeal reference genomes, was used to construct a second maximum likelihood tree (Supplementary Figure 1). Sequences were aligned with MUSCLE, trimmed with trimAL using a 50% gap threshold, and concatenated. The final trimmed and concatenated alignment of 7,178 positions was used for maximum likelihood tree reconstruction using IQ-tree2 v2.0.6, the LG+C60+F+G model, and 1,000 ultrafast bootstraps.

### Group 4 [NiFe]-hydrogenases

Catalytic subunits of group 4 [NiFe]-hydrogenases encoded in Culexarchaeia MAGs were subjected to phylogenetic analysis along with a set of reference sequences extracted from the HydDB (35). Amino acid sequences were aligned using Mafft-LINSi and trimmed using trimAL with a gap threshold of 50%, producing a final alignment of 370 residues. A maximum-likelihood tree was reconstructed using IQ-tree2 v2.0.6 with a best fit model LG+R9 selected according to Bayesian information criterion (BIC), 1,000 ultrafast bootstraps, and option -bnni.

### FtsZ-homologs

Reference amino acid sequences from within the tubulin superfamily (36–38) were downloaded from NCBI (accessed May 2021), combined with sequences obtained from the Culexarchaeia MAGs aligned with Mafft-LINSi, and trimmed using trimAL with a gap threshold of 70%. This produced a final alignment of 313 residues. IQ-tree2 v2.0.6 was used for maximum-likelihood analysis with a best-fit model LG+R5 according to BIC and node support was calculated with 1,000 ultrafast bootstraps.

### Annotation and reconstruction of metabolic potential

Initial analysis of the metabolic potential of the Culexarchaeia MAGs was performed using the annotations provided by the IMG/M database which uses KEGG, COG, pfam, and enzyme ID databases (19). Manual refinement of the IMG/M annotations was done by inspection of gene neighborhoods, and identification of conserved domains and motifs through submission of genes to the NCBI conserved domain database (39) and Interproscan (v5.48) (40). Catalytic subunits of [NiFe]-hydrogenases (COG3261, COG3262, and COG3529) were submitted to the HydDB web portal for classification (35). Putative transmembrane spanning subunits of novel membrane-bound [NiFe]-hydrogenases (Drh, Ehd, Ehe and Ehg complexes) were predicted with TMHMM (41). Supplementary File 3 includes a full list of genes used to construct Figure 3, including the associated KEGG, COG, pfam, Enzyme IDs, and IMG locus tags.

### Assessment of key cellular machinery genes and methanogenesis marker proteins

The Culexarchaeia MAGs and reference genomes from Marsarchaeia, Methanomethylicia, Geoarchaeia, Nezhaarchaeia, *Sulfolobales, Desulfurococcales*, and *Thermoproteales* were screened for the presence and absence of key cellular machinery proteins, identified in previous work (42–44) using arCOG HMMs (45) and HMMER v3.2 (46). A set of previously identified methanogenesis marker proteins (15) were also assessed using arCOGs and HMMER (v3.2) as previously described (15). Following the HMMsearch against Culexarchaeia MAGs and reference archaeal genomes, putative hits (E-value threshold of 1e-3) for each arCOG were manually inspected through BLASTp searches (default settings) against the NCBI non-redundant database and submitted to the NCBI Conserved Domain Database (39). Supplementary File 2 provides the data used to construct Figure 1C and contains the arCOG identifiers used in this analysis along with the presence-absence pattern for each genome.

**Figure 1.**
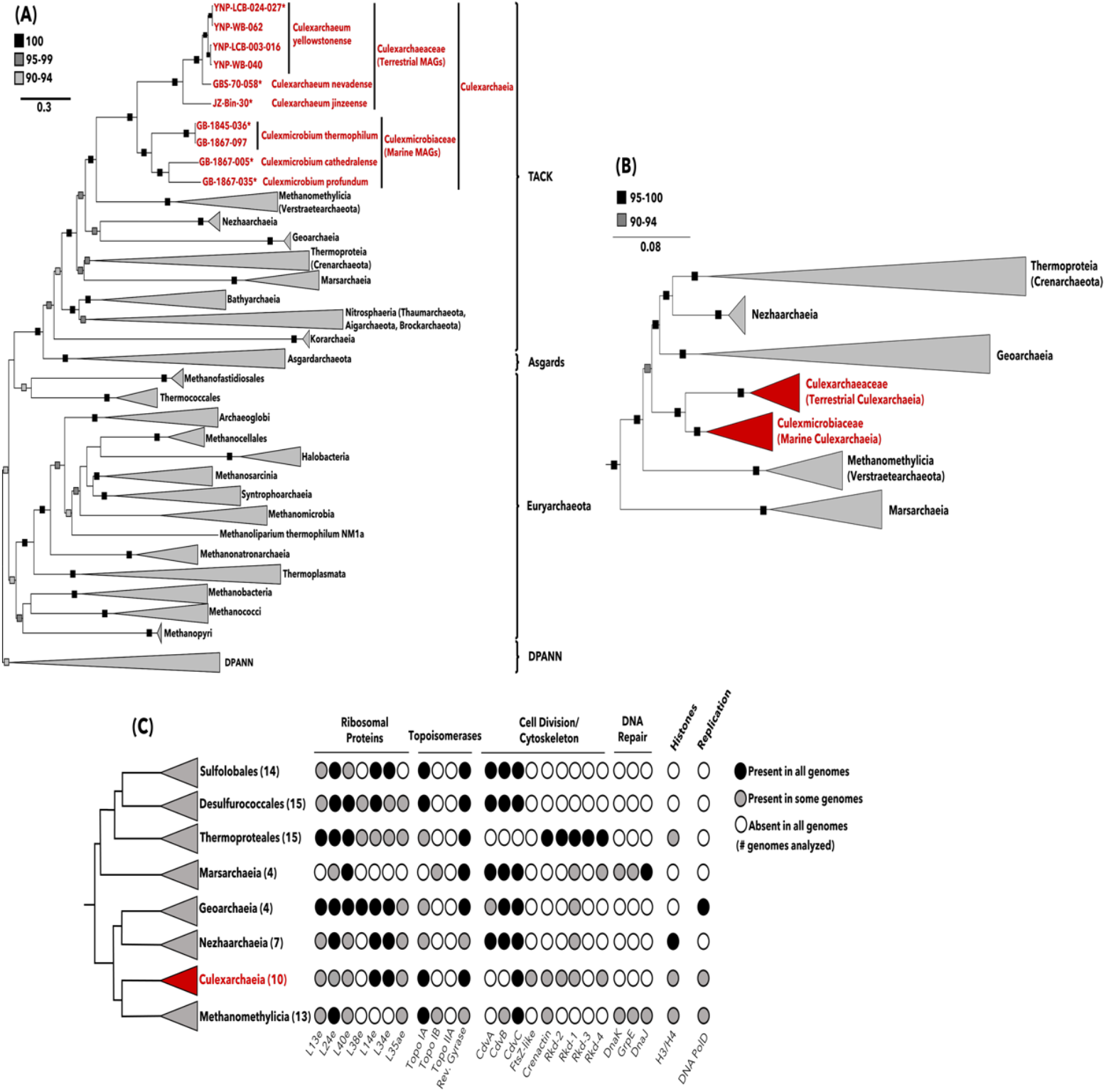
Phylogenetic analysis of Culexarchaeia MAGs and comparison of gene patterns involved in central information-processing machinery. **(A)** Maximum-likelihood tree, inferred with IQtree and the best-fit LG+C60+F+G model, using a concatenated set of 43 conserved arCOGs (Supplementary File 1). Ultrafast bootstrap support values of 100, 95-99, and 90-94 are indicated. *, indicates MAG ID used to designate type species. **(B)** Maximum-likelihood tree, inferred with IQtree and best-fit GTR+F+G model, using 16S rRNA gene sequences longer than 1000 bp. Ultrafast bootstrap support values of 95-100 and 90-94 are indicated. **(C)** Comparison of presence-absence patterns for genes involved in central information-processing machinery among TACK lineages related to *Ca.* Culexarchaeia. Genes found in all genomes analyzed, some genomes analyzed, and absent from all genomes analyzed are indicated with black, gray, and white circles, respectively. The number of genomes screened in each clade are indicated in parentheses, and a full list of the presence-absence pattern for individual genomes can be found in Supplementary File 2.

### Supplementary methodology

Cell extraction protocols and FISH experiments are described in the Supplemental Text.

## Results and Discussion

In this manuscript, we largely refer to MAGs using the recently proposed archaeal Genome Taxonomy Database (GTDB, Release 202) nomenclature (47). However, in the interest of being more accessible in the context of previous work and well-established names in the literature, we also list conventional high-ranking superphylum names (Asgard, DPANN, *Euryarchaeota*, TACK). Parentheses indicate taxa that have undergone significant name changes in the GTDB, but are used interchangeably here (*e.g., Ca.* Methanomethylicia (*Ca.* Verstraetearchaeota); *Thermoproteia* (*Crenarchaeota*)). For simplicity we avoid prefixing candidate taxa with the term “*Ca.*” after they have been introduced.

### Recovery of MAGs

Metagenomic analysis of microbial communities from three hot springs sediment samples in Yellowstone National Park (WY, USA), one sediment sample from Great Boiling Spring (NV, USA) (18), and two deep-sea hydrothermal seep sediment samples from Guaymas Basin (Gulf of California) resulted in the recovery of nine MAGs representative of a new archaeal lineage (Table 1). Additionally, a single MAG (JZ-Bin-30), originally recovered from Jinze Hot Spring (Yunnan, China) (25, 26), that was related to these newly retrieved MAGs, was identified in the Integrated Microbial Genomes (IMG) database. Completeness estimates for the ten MAGs ranged from 89.2-99.0 % and their redundancy ranged from 0-7.79 %. Most of these MAGs contain scaffolds >100 kbp; notably, YNP-LCB-24-027 and GB-1867-005 contain a 1.08 Mbp and 0.84 Mbp scaffold, respectively (Table 1).

**Table 1.** Summary of Culexarchaeia MAGs used in this study. ^a^Estimated with CheckM. ^b^Determined using tRNAscan-SE. ^c^Predicted optimal growth temperature determined using Tome. YNP, Yellowstone National Park; LCB, Lower Culex Basin; WB, Washburn Hot Springs; WY, Wyoming; JZ, Jinze Spring; GBS, Great Boiling Spring; NV, Nevada; GB, Guaymas Basin; N/A, Not available; CDS, Coding sequences; Compl.,

| Genome ID | Origin | Size (Mbp) | Contigs (#) | Largest Scaffold (bp) | CDS | G+C (%) | Compl. (%)<sup>a</sup> | Redundancy (%)<sup>a</sup> | Coverage | Abund. (%)<sup>a</sup> | tRNAs<sup>b</sup> (#) | OGT<sup>c</sup> (°C) |
| --- | --- | --- | --- | --- | --- | --- | --- | --- | --- | --- | --- | --- |
| YNP-LCB-003-016 | Yellowstone National Park, WY, USA | 2.12 | 266 | 105,983 | 2,611 | 38.7 | 94.7 | 0.78 | 72.1 | 0.52 | 37 | 85 |
| YNP-LCB-024-027 | Yellowstone National Park, WY, USA | 1.70 | 42 | 1,082,165 | 1,779 | 39.8 | 97.2 | 0.93 | 55.4 | 0.16 | 47 | 83 |
| YNP-WB-040 | Yellowstone National Park, WY, USA | 1.22 | 141 | 137,472 | 1,515 | 38.9 | 89.7 | 0.93 | 169.6 | 0.67 | 31 | 83 |
| YNP-WB-062 | Yellowstone National Park, WY, USA | 1.63 | 189 | 127,818 | 2,045 | 39.6 | 88.7 | 6.54 | 102.2 | 0.41 | 36 | 87 |
| JZ-Bin-30 | Jinze Hot Spring, Yunnan, China | 1.48 | 33 | 289,617 | 1,588 | 32.4 | 96.7 | 0.00 | 16.2 | N/A | 46 | 82 |
| GBS-70-058 | Great Boiling Spring, NV, USA | 1.42 | 129 | 206,713 | 1,657 | 37.0 | 96.7 | 0.07 | 14.7 | 0.25 | 45 | 83 |
| GB-1845-036 | Guaymas Basin, Mexico | 1.43 | 130 | 112,765 | 1,700 | 37.2 | 99.0 | 3.74 | 17.2 | 0.35 | 37 | 84 |
| GB-1867-005 | Guaymas Basin, Mexico | 1.98 | 25 | 843,365 | 2,245 | 42.2 | 99.0 | 3.74 | 131.7 | 1.86 | 46 | 91 |
| GB-1867-035 | Guaymas Basin, Mexico | 2.04 | 296 | 32,214 | 2,552 | 35.0 | 96.7 | 7.79 | 19.5 | 0.36 | 41 | 89 |
| GB-1867-097 | Guaymas Basin, Mexico | 0.82 | 175 | 18,843 | 1,034 | 38.1 | 89.7 | 2.57 | 13.4 | 0.23 | 33 | 84 |

Completeness; Abund., Abundance; OGT, Optimal growth temperature.

### Phylogenetic placement

Phylogenomic analyses using a concatenated alignment of 43 conserved single copy marker genes (Supplementary File 1) placed the ten MAGs as a monophyletic sister clade to the Methanomethylicia (Verstraetearchaeota) with 100 % bootstrap support (Figure 1A, Supplementary Figure 1). 16S rRNA gene phylogeny further supported placement as a distinct monophyletic clade within the TACK superphylum (100 % bootstrap support; Figure 1B). The average amino acid identity (AAI) values of this lineage compared to other TACK lineages were below 50%, which is comparable to AAI values recovered when comparing Methanomethylicia to other TACK lineages (Supplementary Figure 2A).

Together, these results support the designation of these MAGs as representatives of a distinct archaeal lineage, for which we propose the name “*Candidatus* Culexarchaeia”, after the Culex Basin in Yellowstone National Park. When using the rank-normalized GTDB taxonomy, the phylogenies and AAI comparisons presented here indicate that these MAGs constitute a class-level lineage, Culexarchaeia, separate from Methanomethylicia (Verstraetearchaeota). Alternatively, when using NCBI taxonomy, these MAGs could constitute a phylum-level lineage, “*Candidatus* Culexarchaeota”, separate from Verstraetearchaeota (Methanomethylicia). We support the development of a standardized archaeal taxonomy and from here on refer to this lineage as the class Culexarchaeia. The JZ-Bin-30 MAG identified on IMG was previously proposed to represent a novel species, “*Ca.* Methanomedium jinzeense”, within the order Methanomethylicia (Verstraetearchaeota) (48). However, based on our expanded phylogenetic analysis, we propose to reclassify this MAG as “*Ca.* Culexarchaeum jinzeense” within the Culexarchaeia. Based on phylogenomic analyses, AAI, average nucleotide identity, and pairwise 16S rRNA nucleotide sequence identity analyses, Culexarchaeia can be separated into two families, *Culexarchaeceae* and *Culexmicrobiaceae*, that are composed exclusively of terrestrial and marine representatives, respectively (Figure 1A, Supplementary Figure 2B and 2C). An abbreviated etymology and proposed type material for *Ca.* Culexarchaeia can be found at the end of this manuscript. An extended version can be found in the Supplemental Text.

### Biogeography

To explore the biogeographical distribution of Culexarchaeia, BLASTn searches were used to screen IMG and NCBI non-redundant databases for related 16S rRNA genes (Supplementary File 8). This revealed that Culexarchaeia are globally distributed and are found in circumneutral and slightly acidic pH (5.4-7.8) high temperature (53-83 °C) terrestrial hot spring and deep-sea hydrothermal sediment environments (Figure 2). A small number of 16S rRNA gene sequences were retrieved from deep-sea hydrothermal seep sediment samples in Guaymas Basin with lower temperatures (15 °C; Supplementary File 8). However, the Guaymas hydrothermal system experiences rapidly fluctuating temperature gradients (up to 100 ºC over 50 cm depth) as hydrothermal waters percolate through the sediments (49) (Supplementary Table 2). Consistent with the presence of Culexarchaeia in geothermal environments, the predicted optimal growth temperature based on amino acid composition (29) is above 80 °C for all MAGs with an average of 85 °C (Table 1). Furthermore, all MAGs encode the hyperthermophile-specific enzyme reverse gyrase (Supplementary File 2) that is important for growth at high temperatures (50, 51).

**Figure 2.**
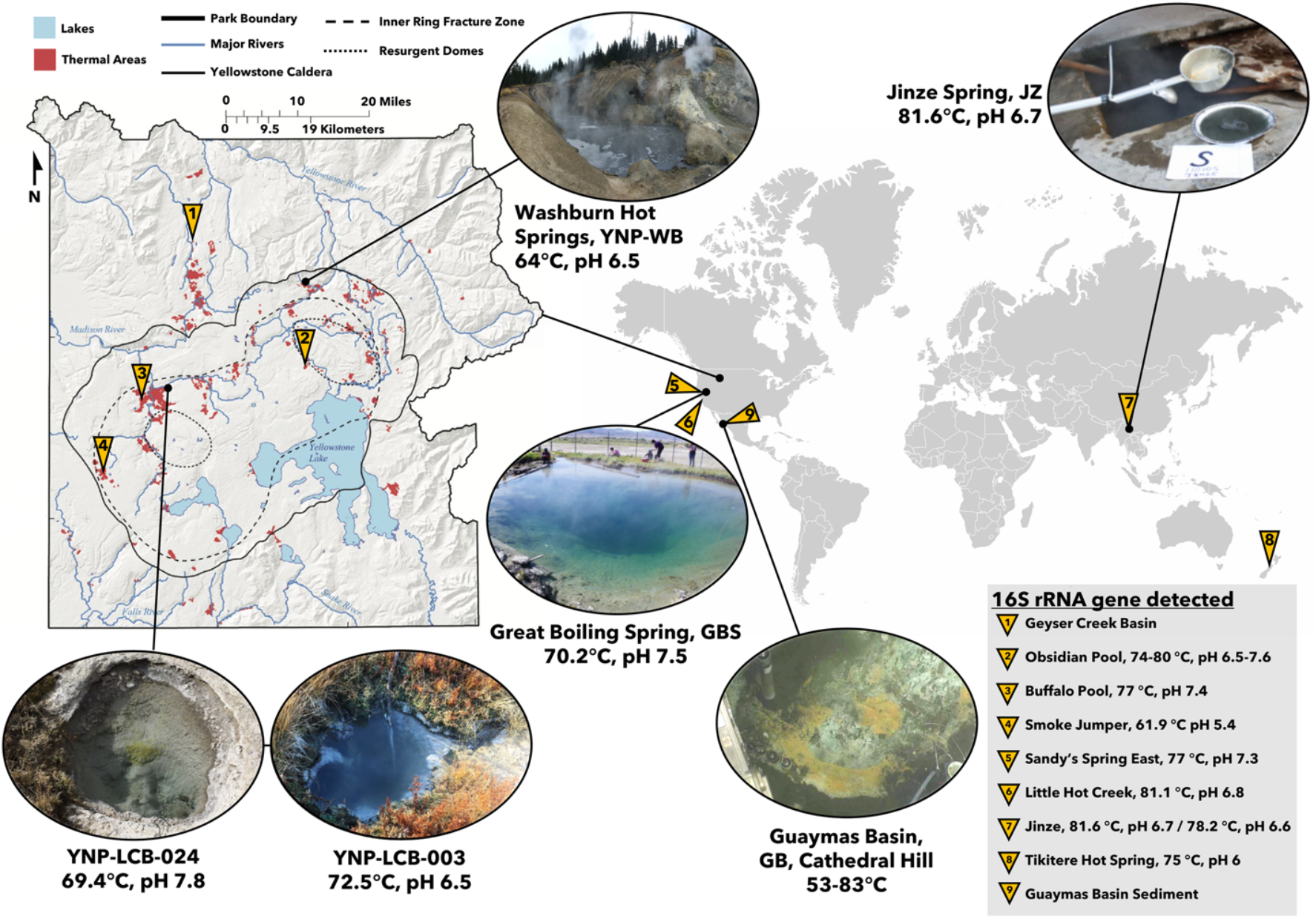
Biogeography of Culexarchaeia. Sites from where Culexarchaeia MAGs were recovered and where 16S rRNA gene sequences have been detected. A full list of samples and associated metadata can be found in Supplementary File 8. Yellowstone map modified from (97), GBS picture modified from (18), and JZ picture modified from (25). Guaymas Basin image courtesy of the Woods Hole Oceanographic Institution.

**Figure 3.**
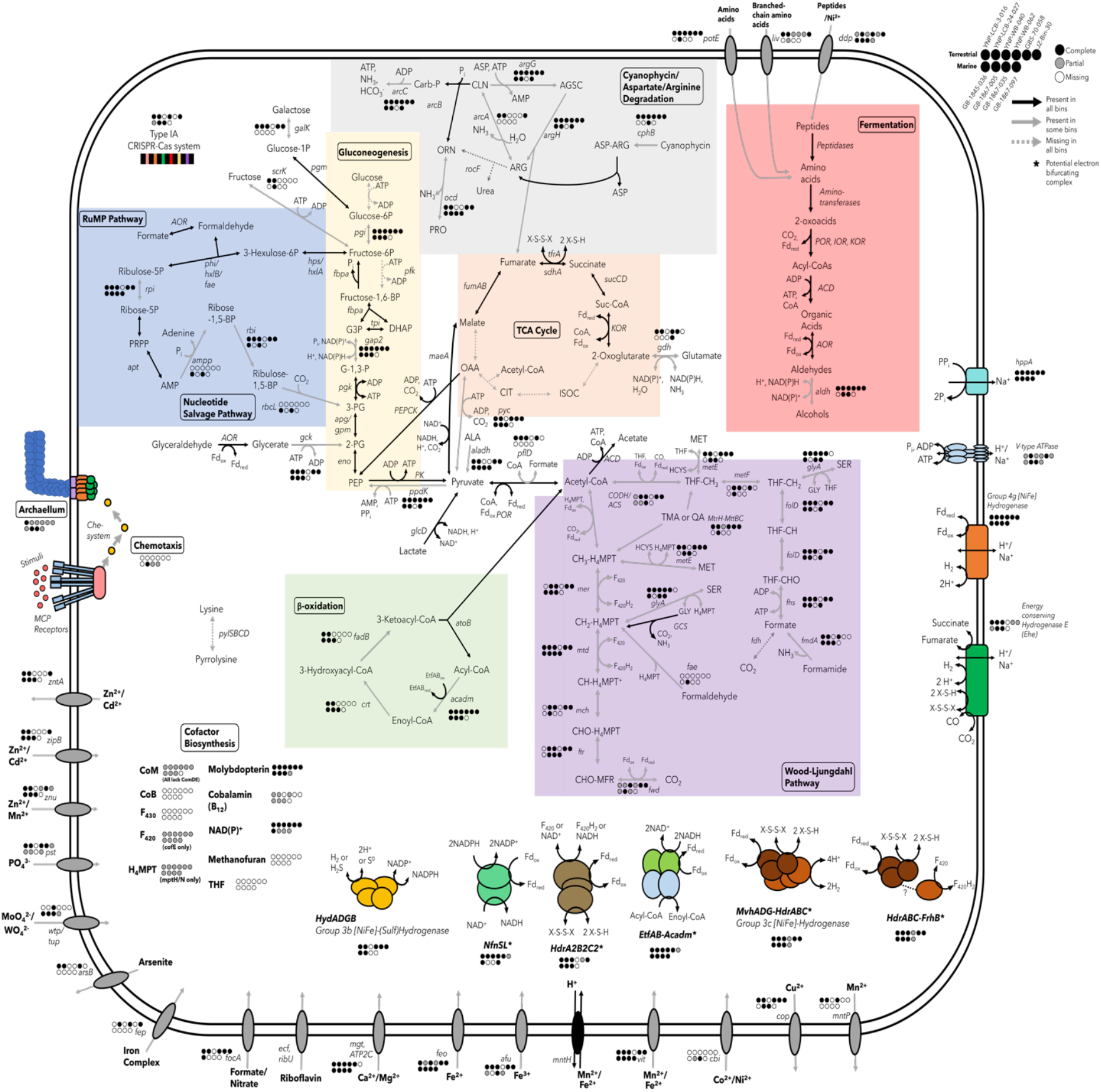
Metabolic potential of Culexarchaeia MAGs. Solid black arrows indicate presence in all bins, solid gray arrows indicate presence in some bins, and dashed gray arrows indicate absence in all bins. For genes encoding multi-subunit complexes and genes present only in some bins (solid gray arrows) - full presence, partial presence, and absence are shown with solid black circles, solid gray circles, and solid white circles respectively. A full list of genes used to construct this figure, gene IDs for each bin, and abbreviations can be found in Supplementary Files 3 and 4. Enzyme complexes with unclear annotation or association are indicated with ‘(?)’. Asterisks (*) indicate potential electron-bifrucating complexes.

We attempted to visualize Culexarchaeia cells in cell extracts from sediment slurries of YNP sites 003 and 024 via catalyzed reporter deposition fluorescence *in situ* hybridization (CARD-FISH) using a newly designed oligonucleotide probe but were unsuccessful. The low relative abundance of Culexarchaeia in YNP sites LCB-003 and LCB-024 (0.52% and 0.16% abundance, respectively; Table 1) may have contributed to our inability to detect them. Alternatively, the failure to detect Culexarchaeia cells could be due to suboptimal probe accessibility, an inability to penetrate the archaeal cell wall under the permeabilization conditions tested, or low ribosome content of relatively inactive cells.

### Comparison of key cellular machinery proteins encoded by Culexarchaeia

To better understand the differences between Culexarchaeia and their closest relatives within the TACK superphylum, we surveyed publicly available genomes within the Geoarchaeia, Marsarchaeia, Methanomethylicia, Nezhaarchaia, and *Thermoproteia* (*Crenarchaeota*), for the presence or absence of proteins involved in key cellular processes (Figure 1C). These marker proteins have been used in previous studies to delineate differences between high-ranking archaeal taxa (42–44) and include archaeal-specific ribosomal proteins, topoisomerases, histones (H3/H4), proteins involved in DNA replication (DNA PolD), DNA repair, cell division, and cytoskeletal proteins. This analysis revealed a lineage-specific presence-absence pattern for Culexarchaeia that differentiates them from other TACK lineages (Figure 1C). In particular, a unique set of cell division and cytoskeletal proteins were found to be encoded in Culexarchaeia MAGs.

### Culexarchaeia encode a unique array of cell division and cytoskeletal proteins

Many lineages in the TACK superphylum (Bathyarchaeia, *Desulfurococcales*, Geoarchaeia, Marsarchaeia Nezhaarchaeia, *Nitrosophaeria* (*Thaumarchaeota*), and *Sulfolobales*) encode a full complement of the ESCRT-III related cell division proteins (52) (CdvABC; Figure 1C). Methanomethylicia, the closest relative to Culexarchaeia, has previously been noted to only encode CdvB and CdvC. This is consistent with our expanded analysis of recently discovered Methanomethylicia representatives (1, 8, 26). In contrast, all Culexarchaeia MAGs only encoded CdvC homologs, which suggests Culexarchaeia may use an alternative mechanism for cell division.

In bacteria, the highly conserved FtsZ tubulin protein mediates the formation of a dynamic division ring during cell division (36). Many archaea (including most members of the *Euryarchaeota*, DPANN, and Asgard, along with a small number of TACK lineages) also encode homologs of the FtsZ tubulin protein, which is critical for cell division (36). Divergent FtsZ homologs encoded by *Haloferax volcanii*, later termed CetZ, were found to not be directly involved with cell division, but instead have roles in maintaining cell shape and motility, which highlights the functional diversity present in this family of proteins (37). Previously, FtsZ homologs had only been noted in other more distantly related TACK lineages including Bathyarchaeia, Caldarchaeles (Aigarchaeota), Korarchaeia, and *Nitrosophaeria* (*Thaumarchaeota*) (52, 53). Most of these TACK lineages each encode at least one homolog that clusters with canonical FtsZ1 and/or FtsZ2 from other archaea, which suggests that these proteins likely play a similar role in cell division. Furthermore, Korarchaeia, Bathyarchaeia, and *Nitrosophaeria* (*Thaumarchaeota*) also encode non-canonical FtsZ-like sequences that have a yet undetermined role in cellular division or cellular shape maintenance (36) (Supplementary Figure 3).

Members of the Culexarchaeia were found to encode a homolog of FtsZ, a protein that is absent in all closely related TACK lineages with the exception of two *Sulfolobales* representatives, which have previously been noted to encode highly divergent and extended proteins belonging to a cluster termed FtsZL1 (38) (Figure 1C, Supplementary Figure 3). Compared to other FtsZ clades, the Culexarchaeia FtsZ-like sequences form a new, deeply branching clade that is distinct from other known tubulin family homologs and branches between the archaeal FtsZ2 group and CetZ group (Supplementary Figure 3). Culexarchaeia FtsZ-like sequences cluster with other uncharacterized sequences from the Thermoplasmata, Bathyarchaeia, Geothermarchaeales, and unclassified Thermoproteia. This novel clade of FtsZ-like proteins represented by Culexarchaeia sequences will require experimental characterization to determine its role in cellular division, cellular shape maintenance, or motility.

In contrast to the Cdv and FtsZ mediated cell division strategies, the cell division and cytoskeletal proteins encoded by the *Thermoproteales* have long been a curiosity because, unlike closely related orders within the *Thermoproteia* (*Crenarchaeota*), they do not encode homologs of the CdvABC or FtsZ proteins (54). Instead, cell division and cellular organization in the *Thermoproteales* have been proposed to rely on homologs of actin (termed crenactin) and several conserved actin-related cytoskeleton proteins (termed arcadins) (54) (Figure 1C). Based on immunostaining of *Pyrobaculum caldifontis*, arcadin-1, arcadin-3, and arcadin-4 were suggested to play auxiliary roles in cell division and cellular shape maintenance, while arcadin-2 was localized in the center of cells between segregated nucleoids, which suggests that it may play a more direct role in cytokinesis (54). Culexarchaeia MAGs recovered from marine environments contained a similar set of proteins (encoding for crenactin, arcadin-1, arcadin-2, and arcadin-4). In contrast, the terrestrial Culexarchaeia MAGs and only encoded arcadin-1, but not crenactin (Supplementary File 2). A few mesophilic members within the Methanomethylicia order Methanomethylicales also encode crenactin, while all other Methanomethylicia genomes analyzed did not encode crenactin (Supplementary File 2). In addition, no arcadin-related proteins were identified in our arCOG survey of Methanomethylicia genomes. A BLASTp search against the non-redundant database using default settings and the Culexarchaeia crenactin sequences as queries revealed that these sequences are more similar to those found in the TACK (*e.g.*, Methanomethylicia and Caldarchaeles) than to those in the Asgard archaea, which are more similar to eukaryotic actin sequences (55).

The set of cell division, cell shape, and cytoskeleton proteins encoded by the Culexarchaeia suggests that they have a unique cell division and maintenance process. Similarly, members of the Caldarchaeles (Aigarchaeota) have been noted to encode a unique complement of cell division (FtsZ2, CdvBC) and cytoskeletal proteins (Crenactin, arcadin-1, arcadin-2) (56). Likewise, some Asgard archaea (*e.g.,* Heimdallarchaeota, Odinarchaeota) have been found to encode a mixture of proteins possibly involved in cell division machinery (CdvBC in addition to FtsZ1 and/or FtsZ2) and actin-like sequences that are closely related to eukaryotic homologs (1, 52). It is yet unknown if, or how, these diverse systems interact with each other during the cell cycle (43, 56). All together, these data suggest that Culexarchaeia, along with other TACK and Asgard lineages, are critical to understanding the diversity and evolution of cell division and cytoskeletal proteins within the *Archaea*.

### Metabolic potential of Culexarchaeia

Culexarchaeia have the potential to live a generalist lifestyle, with the ability to metabolize a variety of substrates including proteins, amino acids, fatty acids, sugars, and methyl compounds (Figure 3; Supplementary File 3).

### Peptide degradation and fermentation

Culexarchaeia encode peptide and amino acid transporters that can import exogenous peptides or amino acids that can be catabolized to their respective 2-oxoacids via encoded peptidases and amino-transferases (57, 58). Ferredoxin oxidoreductases (POR, IOR, KOR) can then convert these 2-oxoacids to their respective Acyl-CoA substrate and reduce a low-potential ferredoxin (58, 59). Culexarchaeia encode multiple Acyl-CoA synthetases that can convert the Acyl-CoA substrates to organic acids and generate ATP via substrate level phosphorylation (58, 59). Culexarchaeia also encode multiple aldehyde-ferredoxin oxidoreductases (AOR) and alcohol dehydrogenases that could act to interconvert organic acids to aldehydes and aldehydes to alcohols, respectively (59–61). This would remove excess reducing equivalents (NAD(P)H and/or reduced ferredoxin) produced during fermentation. Alternatively, or in addition to, reduced ferredoxins produced by ferredoxin oxidoreductases could also be oxidized by a membrane bound group 4g [NiFe]-hydrogenase concomitantly with H2 production and the transport of ions outside of the cell (Figure 3). This scenario would be akin to the metabolism of *Pyrococcus furiosus*, which uses a membrane bound hydrogenase and ATP synthase to conserve energy during hydrogenogenic fermentation (62). Additional fermentative capacity is possible through the conversion of pyruvate to lactate or alanine through the activity of lactate dehydrogenase (GlcD) or alanine dehydrogenase (Aladh), respectively.

### Cyanophycin degradation

All terrestrial MAGs and one marine MAG encode cyanophycinase, a peptidase capable of degrading cyanophycin, a carbon and nitrogen storage polymer (63). Cyanophycin is commonly found in cyanobacteria and some heterotrophic bacteria and is composed of repeated aspartate and arginine dipeptides (64). After degrading cyanophycin polymers, Culexarchaeia could further metabolize arginine via the arginine deiminase pathway (65) (*arcA, arcB, arcC*; Figure 3) to generate ATP, while aspartate could be used in central metabolism via aspartate aminotransferase, converting it to the TCA intermediate oxaloacetate. To the best of our knowledge, cyanophycin degradation by archaea has had extremely limited discussion in the literature and no experimental evidence of archaeal cyanophycin degradation has yet been obtained. Additionally, BLASTp searches against the non-redundant NCBI database with default settings using the Culexarchaeia cyanophycinase as a query suggests that a small assortment of Euryarchaea and Asgardarchaea also encode potential cyanophycinase homologs and that the role of archaea in cyanophycin cycling is currently underappreciated (66). Given the apparent limited capacity for other archaea to potentially degrade this substrate, we speculate that this substrate could be used in combination with antibiotics to aid in isolating Culexarchaeia, because cyanophycin could act as a sole carbon, nitrogen, and energy source.

### Wood-Ljungdahl pathway and methylotrophy

Culexarchaeia encode portions of both the tetrahydrofolate (THF) and tetrahydromethanopterin (H4MPT) methyl branches of the Wood-Ljungdahl pathway (WLP) (Figure 3). None of the 38 marker genes previously identified in methanogens, including methyl coenzyme-M reductase (MCR) and the biosynthesis genes for the MCR cofactor F430 (15), were identified in any Culexarchaeia MAGs (Supplementary File 6). Furthermore, Culexarchaeia only encode the MtrH subunit of the tetrahydromethanopterin S-methyltransferase (MTR) complex that is involved in the transfer of a methyl group to Coenzyme-M, which forms methyl-CoM during hydrogenotrophic or methylotrophic methanogenesis (67). These *mtrH* genes were co-located with genes encoding for trimethylamine methyltransferases (*mttB*; COG5598), corrinoid proteins (COG5012), and a corrinoid activation protein (COG3984) (Supplementary Figure 4). It was recently shown that non-pyrrolysine members of the MttB superfamily, which exhibit similar co-localization of genes (*mtrH*-*mttB*-*mttC*), are involved in the demethylation and degradation of quaternary amines such as glycine betaine or *L*-carnitine (68, 69). Culexarchaeia lack the genes necessary for biosynthesis of pyrrolysine (*pylSBCD*), which supports the hypothesis that this complex could be involved in the catabolism of quaternary amine compounds or yet unknown methylated substrates. Methyl groups transferred by methyltransferase complexes could potentially enter either the THF or H4MPT branches of the WLP, producing THF-CH3 or CH3-H4MPT, respectively (Figure 3). Furthermore, Culexarchaeia may compete for methylated compounds with other methylotrophic organisms found in geothermal environments, including methylotrophic methanogens, methylotrophic bacteria, or the recently identified Brockarchaeota (9).

Together, these findings suggest that Culexarchaeia are incapable of methanogenesis, and that the WLP would confer metabolic flexibility by acting as a major hub for the entry and exit of multiple substrates. Specifically, if operating in the reductive direction, acetogenesis could occur from CO_2_, formate, or methylated compounds (potentially trimethylamine or quaternary amines) (Figure 3). Formate could be produced through the activity of pyruvate-formate lyase (PflD), formamidase (FmdA), or through interconversion of formaldehyde and formate by aldehyde-ferredoxin oxidoreductases (AOR; Figure 3). Alternatively, the WLP could be used in the oxidative direction to oxidize Acetyl-CoA or methylated compounds to CO_2_ or formate. Culexarchaeia may also use the WLP for protein degradation because they encode serine hydroxymethyltransferase, which can produce a methylene-H_4_MPT and glycine. The glycine cleavage system (GCS) could then convert glycine to methylene-H_4_MPT, CO_2_, and NH_3_ (70) (Figure 3).

### Beta-oxidation

Culexarchaeia encode genes for the metabolism of fatty acids through beta-oxidation. This includes medium chain Acyl-CoA dehydrogenases (Acadm), electron transfer flavoproteins (EtfAB), Enoyl-CoA hydratase (Crt), 3-hydroxybutyryl-CoA dehydrogenase (FadB), and Acetyl-CoA C-acetyltransferase (AtoB) (71). Acetyl-CoA produced by this pathway could then be metabolized by oxidation through the WLP or by Acyl-CoA synthetases to produce ATP through substrate level phosphorylation (Figure 3). Additionally, electron transfer flavoproteins and Acyl-CoA dehydrogenase may form an electron-bifurcating complex (EtfAB-Acadm), as was previously suggested for fatty acid-degrading *Firmicutes* (72).

### Carboxydotrophy

Culexarchaeia MAGs encode putative aerobic carbon-monoxide dehydrogenases (CoxLMS), which suggests a capacity for carboxydotrophy through this enzyme complex. Culexarchaeia also encode genes involved in molybdopterin cofactor biosynthesis (Figure 3), which could be involved with such a molybdoenzyme. However, consistent with an analysis of Cox proteins in Caldarchaeles (Aigarchaeota) and a broader analysis of form I Cox proteins in bacteria and archaea, all Culexarchaeia CoxL homologs lack conserved active site residues (VAYRCSFR) found in other characterized Cox proteins and do not cluster with other characterized form I Cox proteins. Together, this suggests these enzymes may use an alternative substrate (73–75). However, carboxydotrophic potential in Culexarchaeia may exist via a [NiFe]-hydrogenase complex that interacts with carbon monoxide dehydrogenase subunit A (CdhA; discussed below).

### Gluconeogenesis, RuMP Pathway, TCA cycle, and Nucleotide Salvage Pathway

Culexarchaeia encode important enzymes involved in gluconeogenesis, including phosphoenolpyruvate carboxykinase (PEPCK) and pyruvate:phosphate dikinase (PpdK) for the production of phosphoenolpyruvate (PEP) from oxaloacetate and pyruvate, respectively (76). While they lack a gene for phosphofructokinase, all MAGs encode fructose bisphosphate aldolase/phosphatase (Fbpa) (77). Together, this suggests Culexarchaeia are capable of gluconeogenesis (76).

Culexarchaeia have the potential to metabolize a variety of sugars and aldehydes. Several MAGs encode the proteins necessary for using fructose or galactose, which could be converted to fructose-6-phosphate for entry into the ribulose monophosphate (RuMP) pathway (Figure 3). The RuMP pathway could be used for formaldehyde assimilation/detoxification or biosynthesis of nucleotides (78). Alternatively, formaldehyde produced by 6-phospho-3-hexuloisomerase (HxlB) could be converted to formate via an aldehyde:ferredoxin oxidoreductase and then enter the THF branch of the WLP via formate-tetrahydrofolate ligase (Fhs). A single marine MAG, GB-1867-005, encoded a formaldehyde-activating enzyme (Fae) that could incorporate formaldehyde into the H4MPT branch of the WLP. Glyceraldehyde may be scavenged through conversion to glycerate by an aldehyde-ferredoxin oxidoreductases and glycerate could subsequently be converted to 2-phosphoglycerate by glycerate kinase (Gck) (Figure 3).

Culexarchaeia encode an incomplete tricarboxylic acid (TCA) cycle that may function as a hub for protein fermentation and conversion of TCA intermediates (malate and oxaloacetate) to pyruvate for gluconeogenesis or conversion to Acetyl-CoA (Figure 3). Peptide degradation could produce 2-oxoglutarate and glutamate through the activity of aminotransferases and peptidases, respectively. Then, glutamate dehydrogenase (Gdh) could facilitate the interconversion of these substrates. 2-oxoglutarate could be metabolized to produce reduced ferredoxin and succinyl-CoA via 2-oxoglutarate:ferredoxin oxidoreductase (KOR). Succinyl-CoA could then be stepwise converted to succinate, fumarate, and malate through the activities of succinyl-CoA synthetase (SucCD), fumarate reductase (TfrA), and fumarate hydratase (FumAB), respectively (Figure 3). Malate could be converted to pyruvate by the decarboxylating malic enzyme (MaeA) and enter gluconeogenesis through the conversion to PEP by pyruvate:phosphate dikinase (Ppdk). Similar to Methanomethylicia, two marine MAGs encode a full nucleotide salvage pathway that includes Ribulose 1,5-bisphosphate carboxylase (RbcL) for conversion of adenosine monophosphate (AMP) to 3-phosphoglycerate (3-PG), which could be metabolized to acetyl-CoA (12).

### Extensive capacity for H_2_-cycling and many potential electron bifurcating/confurcating complexes

The large number of hydrogenases and putative electron-bifurcating complexes encoded by Culexarchaeia MAGs suggest that these enzymes are important in their metabolism. Flavin-based electron bifurcation involves the coupling of an exergonic redox reaction to drive an endergonic redox reaction (79). Electron bifurcation has been found to be important in balancing redox carriers within the cell and increasing efficiency of cell metabolism, particularly for anaerobic organisms that grow on low energy-yielding substrates (80, 81). Culexarchaeia encode five different potential electron bifurcating complexes, which places them among a small number of genomes found to encode more than three bifurcating complexes. These enzymes may provide advantages in energy efficiency and competition over low energy-yielding substrates *in situ* (79–81).

Culexarchaeia encode multiple group 3 and 4 [NiFe]-hydrogenases, which indicates that they could either produce or consume H2. The group 3c [NiFe]-hydrogenase (MvhADG) may form a bifurcating complex with HdrABC, as found in other diverse archaea, which allows for the oxidation of H2 to be coupled to the reduction of ferredoxin and a disulfide compound (82). Alternatively, given the lack of FrhAG in Culexarchaeia MAGs, HdrABC could form a confurcating complex with FrhB, thus coupling the oxidation of a disulfide and ferredoxin to the reduction of F_420_, similar to a scenario proposed for *Methanoperedens* that don’t encode FrhAG (83). Culexarchaeia also encode three other complexes with the potential to perform electron bifurcation: HdrA2B2C2 (discussed below), EtfAB-Acadm (discussed above with beta oxidation), and NADH-dependent ferredoxin:NADP^+^ oxidoreductase (NfnSL; Figure 3), By coupling the oxidation of NAD(P)H with the reduction of NAD(P)^+^, this enzyme could play a key role in cellular redox balancing (84).

Culexarchaeia encode a group 3b [NiFe]-hydrogenase that may couple the reversible oxidation of NADPH to the reduction of protons to evolve H2 (85). Characterization of the group 3b [NiFe]-hydrogenase in *Pyrococcus furiosus* revealed that it had sulfhydrogenase activity. This would allow the cell to couple the oxidation of NADPH with the reduction of elemental sulfur (S^0^) or polysulfide to H2S (85, 86). Thus, Culexarchaeia may be involved in sulfur cycling through the activity of this potential sulfhydrogenase. Reduced ferredoxin produced through fermentative metabolism could be oxidized by a membrane-bound group 4g [NiFe]-hydrogenase, coupled to the translocation of ions and the production of H2 (Figure 3). Culexarchaeia also encode a novel group 4 [NiFe]-hydrogenase (termed Ehe, discussed below) that may interact with disulfides, CO, or succinate. Furthermore, the production of H2 (or formate) by Culexarchaeia could potentially form the basis of syntrophic interactions with other community members through the exchange of these compounds.

### An expanded diversity of potential HdrB-interacting hydrogenases

A recent evaluation of MAGs affiliated with the Methanomethylicia, the closest relatives to Culexarchaeia, revealed genes encoding a novel membrane bound [NiFe]-hydrogenase complex (termed Ehd) that were co-located with genes encoding heterodisulfide reductase subunits (HdrBC) and an ion antiporter subunit (15) (Figure 4B, 4F). This Ehd complex was suggested as an alternative pathway to couple the oxidation of H2 to the reduction of the heterodisulfide CoM-S-S-CoB and translocation of ions outside the cell, which would allow for energy conservation during methylotrophic methanogenesis (15). A similar membrane-bound hydrogenase complex (termed Ehe here) is encoded in Culexarchaeia MAGs (Figure 4A, 4E). The Ehd and Ehe gene clusters have the same core structure containing genes for a [NiFe]-hydrogenase complex, HdrBC, and an ion antiporter subunit. However, the genes encoding the Ehe complex are co-located with genes encoding succinate dehydrogenase subunit A (SdhA), carbon monoxide dehydrogenase subunit A (CdhA), a predicted bifurcating heterodisulfide reductase complex (HdrA2B2C2) (87), and a glycine cleavage protein H (GcvH). This suggests that carbon monoxide or succinate oxidation could be coupled to the reduction of protons and the translocation of ions to the outside of the cell. Thermodynamics of succinate oxidation coupled to H2 production indicate that this reaction becomes favorable at temperatures >65 °C (88), which aligns with some of the high-temperature environments that Culexarchaeia inhabit (50-83ºC). Additionally, CO is a common trace gas in geothermal systems and could potentially be oxidized by this complex and coupled to the reduction of protons to H2 to fuel carboxydotrophic growth (89).

**Figure 4.**
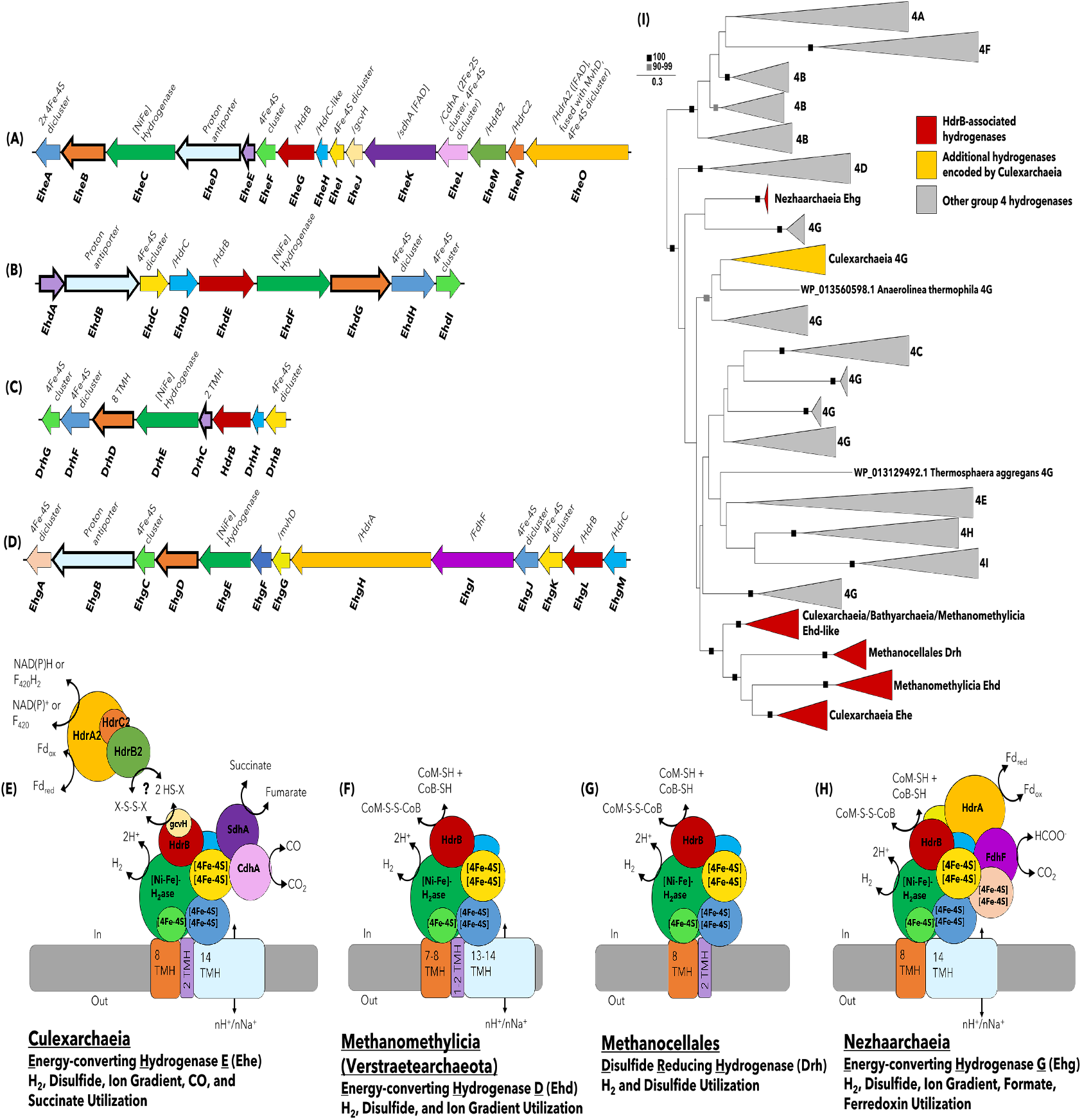
Modular HdrB-interacting [NiFe] Hydrogenases present in Culexarchaeia, Methanomethylicia (Verstraetearchaeota), Methanocellales, and Nezhaarchaeia. **(A,E)** Gene structure and tentative model of the Ehe complex proposed for Culexarchaeia, **(B,F)** Ehd complex proposed for Methanomethylicia (Verstraetearchaeota) (modified after (70)), **(C,G)** Drh complex proposed for Methanocellales, and **(D,H)** Ehg complex proposed for Nezhaarchaeia. Transmembrane helices (TMH) were predicted using TMHMM and sequences in panels A-D with TMHs are indicated with a thick border. Sequences that share a similar structural/functional role in these complexes are indicated with the same color. **(I)** Maximum-likelihood phylogenetic reconstruction of the catalytic subunit of group 4 [NiFe]-hydrogenases. Ehe, Ehd, Drh, and Ehg clades are displayed in red. Additional Culexarchaeia sequences belonging to group 4 [NiFe]-hydrogenases are labeled in yellow.

Alternatively, H2 oxidation could also be coupled to the reduction of an unknown disulfide (via HdrB) that could be generated through the activity of the HdrA2B2C2 or HdrABC-MvhADG bifurcating complexes. The identity of this disulfide is unknown, but Culexarchaeia lack a full biosynthetic pathway for CoM and CoB, which suggests that an alternative disulfide may be used. The GcvH protein, encoded within the Ehe gene cluster, typically functions in the glycine cleavage pathway to coordinate a disulfide containing lipoyl moiety (90, 91). This suggests GcvH could play a similar role in coordinating a disulfide containing moiety during potential interactions with HdrB and/or the HdrA2B2C2 bifurcating complex. Determining directionality of the reaction catalyzed by this complex (oxidation or reduction of H2) and how the diverse set of associated modules (HdrB, CdhA, SdhA, GcvH, and ion transporter) are regulated will require experimental work.

Searches using the genes encoding HdrB and [NiFe]-hydrogenase catalytic subunit revealed gene clusters with similar synteny to the ones encoding the Ehd and Ehe complexes in the methanogenic *Methanocellales* (Figure 4C, 4G; denoted Drh) and in the predicted methanogenic Nezhaarchaeia (Figure 4D, 4H; termed here Ehg) (8, 92). Both gene clusters that encode the Drh and Ehg complexes retain the core structure of the [NiFe]-hydrogenase and HdrB genes, but co-located genes differ. Specifically, the Drh complex in *Methanocellales* lacks an ion antiporter subunit, which suggests that it is not involved in energy conservation via the production of an ion gradient, but instead could be directly involved in the reduction of CoM-S-S-CoB coupled to H2 oxidation (92). The proposed Ehg gene cluster in Nezhaarchaeia MAGs is co-located with genes for a formate dehydrogenase subunit (FdhF), HdrA subunit, MvhD subunit, and an ion antiporter subunit. This suggests that oxidation of H2, formate, or ferredoxin might be coupled to the direct reduction of CoM-S-S-CoB and the translocation of protons outside of the cell, thus conserving energy during their proposed hydrogenotrophic methanogenesis.

Phylogenetic analysis of the [NiFe]-hydrogenase catalytic subunit revealed that the Ehd, Ehe, and Drh complexes form a distinct cluster relative to other classified group 4 [NiFe]-hydrogenases. In contrast, the Ehg complex in Nezhaarchaeia forms a separate clade near a set of group 4g [NiFe]-hydrogenases (Figure 4I). Additionally, JZ-Bin-30 encodes a second putative HdrB and membrane associated [NiFe]-hydrogenase complex that clusters with Bathyarchaeia and Methanomethylicia sequences (Figure 4I). This cluster was termed Ehd-like, due to the similarity in synteny to the Ehd complex in Methanomethylicia, which lacks associated SdhA, CdhA, and GcvH subunits.

Importantly, these observations suggest that the strategy of coupling disulfide reductase activity to membrane bound [NiFe]-hydrogenases and ion translocation may be common in both methanogenic and non-methanogenic archaeal lineages. The apparent modular nature of these complexes may have allowed for diversification in substrate usage in the case of Culexarchaeia (succinate, CO) and Nezhaarchaeia (formate) for their respective metabolisms. Culturing representatives from the Culexarchaeia, Methanomethylicia, or Nezhaarchaeia lineages would enable investigations into the function of these disulfide-interacting group 4 [NiFe]-hydrogenases in the metabolism of these organisms.

## Conclusion

Here we report on a new archaeal class within the TACK superphylum, “*Candidatus* Culexarchaeia”, that is most closely related to the proposed methylotrophic and methanogenic lineage “*Ca.* Methanomethylicia” (Verstraetearchaeota) (12). In contrast to their closest relatives, Culexarchaeia encode the potential for non-methanogenic anaerobic methylotrophy but not methanogenesis. Furthermore, they encode the potential for a generalist lifestyle, with the capacity to use a diverse set of organic (sugars, lipids, proteins) and inorganic (H_2_, CO, S^0^) substrates. Notably, the potential for anaerobic non-methanogenic methylotrophy and cyanophycin degradation, which to our current knowledge are not widespread among archaea, suggests that Culexarchaeia may be important in cycling these compounds in extreme environments and that these putative functions could be exploited in future cultivation attempts. The biogeographic distribution of terrestrial and marine Culexarchaeia indicates that they are found in high temperature (>50° C) and circumneutral to slightly acidic (pH 5.4-7.8) environments. The capacity to metabolize many organic and inorganic substrates suggests that they could be important in the biogeochemical cycles of hydrogen, carbon, and sulfur within these diverse geothermal habitats, even at low relative abundances. The metabolic versatility encoded by Culexarchaeia could imply adaptations to changing nutrient conditions present in dynamic geothermal systems, because they could access a wide variety of compounds for their carbon and energy needs. Additionally, Culexarchaeia expand the diversity of cell division and cytoskeletal proteins in the TACK archaea, as they encode FtsZ-like, CdvBC, crenactin, and arcadin proteins. Future studies will need to determine how these diverse cell division and cytoskeletal proteins contribute to cell cycle and maintenance processes in Culexarchaeia, and could aid in understanding other lineages that similarly encode multiple division and cytoskeletal systems (*e.g.*, Caldarchaeles (Aigarchaeota), Heimdallarchaeota, and Odinarchaeota).

The phylogenetic placement of Culexarchaeia aids in reconstructing the evolutionary history of TACK archaea. Particularly, their relatedness to the proposed methanogenic Methanomethylicia and Nezhaarchaeia, may provide insights into the evolutionary transitions between methanogenic and non-methanogenic lineages. Currently, the evolution of the WLP as well as the MCR and MTR complexes in the TACK archaea are not well understood, which precludes our understanding of the evolutionary history of methane metabolism and whether the last ancestral methanogen used methyl compounds or H_2_/CO_2_ (15, 48, 93). Culexarchaeia encode the potential for anaerobic non-methanogenic methylotrophy via the WLP, but do not encode MCR or MTR, which provides a stark contrast between the methyl-utilizing Methanomethylicia (which encode MCR, but don’t encode the WLP and MTR) and the H_2_/CO_2_ utilizing Nezhaarchaeia (which encode the WLP, MCR, and MTR). If methanogenesis was the metabolism present in the last common ancestor of these lineages, Culexarchaeia will aid in understanding shifts from methanogenic to non-methanogenic lifestyles in the TACK archaea. This type of transition is considered to have been an important process in archaeal evolution and is partially responsible for the observed patchwork distribution of MCR-encoding lineages (15, 94, 95). We expect that as more deeply branching lineages in the TACK superphylum are recovered, this will help resolve the complicated history of these metabolic transitions and further our understanding of the metabolic capabilities encoded by this diverse superphylum.

### Etymology and proposal of type material

Following the recommendations by Chuvochina *et al.* 2019 (96), we assign type genomes and propose the following provisional taxonomic assignments. For a tabular list see Supplementary Table 2.

Candidatus Culexarchaeia class nov.

Cu.lex.ar.chae’ia. N.L. neut. n. Culexarchaeales, type order of the class; L. -ia, ending to designate a class; N.L. fem. pl. n. Culexarchaeia, the Culexarchaeum class. The description is the same as for Candidatus Culexarchaeum gen. nov.

Candidatus Culexarchaeales order nov.

Cu.lex.ar.chae’ales. N.L. neut. n. Culexarchaeaceae, type family of the order; L. -ales, ending to designate an order; N.L. fem. pl. n. Culexarchaeales, the Culexarchaeum order. The description is the same as for Candidatus Culexarchaeum gen. nov.

Candidatus Culexarchaeaceae fam. nov.

Cu.lex.ar.chae.ace’ae. N.L. neut. n. Culexarchaeum, type genus of the family; L. -aceae ending to designate a family; N.L. fem. pl. n. Culexarchaeaceae, the Culexarchaeum family. The description is the same as for Candidatus Culexarchaeum gen. nov.

Candidatus Culexarchaeum gen. nov.

Cu.lex.ar.chae’um N.L. neut. n. Culex, referring to the Culex Basin; N.L. neut. n. archaeum, an archaeon; N.L. neut. n. Culexarchaeum, archaeon of Culex, referring to the Culex Basin of Yellowstone National Park, where this lineage was discovered. The type species is Candidatus Culexarchaeum yellowstonense.

Candidatus Culexarchaeum yellowstonense sp. nov.

yel.low.ston.en’se N.L. neut. adj. yellowstonense, from Yellowstone National Park. This uncultured lineage is represented by bin YNP-LCB-024-027, recovered from an unnamed hot spring in the Culex Basin of Yellowstone National Park. The bin has an estimated completeness of 97.2% and a contamination of 0.93% and contains 16S rRNA, 23S rRNA and 5S rRNA genes.

Candidatus Culexarchaeum nevadense sp. nov.

ne.va.den’se N.L. neut. adj. nevadense, from Nevada. This uncultured lineage is represented by bin GBS-70-058, recovered from Great Boiling Springs in Nevada. The bin has an estimated completeness of 96.7% and a contamination of 0.07% and contains 16S rRNA, 23S rRNA, and 5S rRNA genes.

Candidatus Culexarchaeum jinzeense sp. nov.

jin.ze.en’se N.L. neut. adj. jinzeense, from Jinze. Represented by bin JZ-bin-30, recovered from a geothermal well in Jinze, Yunnan Province, China). The bin has an estimated completeness of 96.7%, a contamination of 0%, and contains 16S rRNA, 23S rRNA, and 5S rRNA genes. For a justification for the reclassification of this previously published bin (48), please see Supplemental Text.

Candidatus Culexmicrobiaceae fam. nov.

Cu.lex.mi.cro.bi.ace’ae. N.L. neut. n. Culexmicrobium, type genus of the family; L. -aceae ending to designate a family; N.L. fem. pl. n. Culexmicrobiaceae, the Culexmicrobium family. The description is the same as for Candidatus Culexmicrobium gen. nov.

Candidatus Culexmicrobium gen. nov.

Cu.lex.mi.cro.bi.ace’ae. N.L. neut. n. Culex, referring to the Culex Basin; N.L. neut. n. microbium, a microbe; N.L. neut. n. Culexmicrobium, microbe of Culex, referring to the Culex Basin of Yellowstone National Park, where this lineage was discovered. The type species is Candidatus Culexmicrobium cathedralensis.

Candidatus Culexmicrobium cathedralense sp. nov.

ca.the.dra.len’se N.L. neut. adj. cathedralense, from Cathedral Hill, a deep-sea hydrothermal vent in Guaymas Basin. This uncultured lineage is represented by bin GB-1867-05, recovered from deep-sea hydrothermal sediment in Guaymas Basin in the Gulf of California (i.e., Sea of Cortés). The bin has an estimated completeness of 99%, a contamination of 3.74%, and contains 16S rRNA, 23S rRNA and 5S rRNA genes.

Candidatus Culexmicrobium thermophilum sp. nov.

ther.mo.phil’um N.L. neut. adj. thermophilum, heat-loving. This uncultured lineage is represented by bin GB-1845-036, recovered from deep-sea hydrothermal sediment in Guaymas Basin in the Gulf of California (i.e., Sea of Cortés). The bin has an estimated completeness of 99%, a contamination of 3.74%, and contains 16S rRNA, 23S rRNA and 5S rRNA genes.

Candidatus Culexmicrobium profundum sp. nov.

pro.fund’um N.L. neut. adj. profundum, deep. Represented by bin GB-1867-035, recovered from deep-sea hydrothermal sediment in Guaymas Basin in the Gulf of California (i.e., Sea of Cortés). The bin has an estimated completeness of 96.7%, a contamination of 7.79%, and contains a 5S rRNA gene.

## Author contributions

A.J.K, Z.J.J., and R.H. designed the research. M.L. and R.H. collected molecular and geochemical samples from YNP hot springs. A.J.K. and Z.J.J. processed metagenome data, evaluated the MAGs, and performed phylogenomic and 16S rRNA analyses. A.J.K. reconstructed the metabolic potential of Culexarchaeia, performed the phylogenetic analyses of individual proteins, and ran FISH experiments. V.K. provided useful discussion and contributed to assessing metabolic potential. A.J.K. and R.H. wrote the manuscript, which was edited by all authors.

## Acknowledgements

We thank Drs. Barbara MacGregor, William Inskeep and Mensur Dlakić, as well as Brian Hedlund and Ramunas Stepanauskas for permitting use of their metagenomes and for providing metadata information for Guaymas Basin sediments, Washburn Hot Springs, and Great Boiling Spring, respectively. We acknowledge Christine Gobrogge at Montana State University’s Environmental Analytical Laboratory for assisting with geochemical analyses. We further thank Drs. Brian Hedlund and Marike Palmer for discussing nomenclatural types and taxonomic naming and Grayson Chadwick for helpful comments on the manuscript. This study was funded through a NASA Exobiology program award to R.H. (80NSSC19K1633). A.K. and M.L. were supported in part by the Thermal Biology Institute and Montana State University’s Vice President’s Office of Research, Economic Development and Graduate Education. V.K. was supported in part by a grant from the W.M. Keck Foundation. A portion of this research was performed under the Facilities Integrating Collaborations for User Science (FICUS) program (proposal: 10.46936/fics.proj.2017.49972/6000002) and used resources at the DOE Joint Genome Institute (https://ror.org/04xm1d337), which is a DOE Office of Science User Facility operated under Contract No. DE-AC02-05CH11231. We thank the US National Park Service, in particular Annie Carlson at the Yellowstone Center for Resources, for permitting work in YNP under permit number YELL-SCI-8010.

## Data Availability

The raw metagenomes used to recover MAGs of Culexarchaeia representatives are publicly available through the JGI IMG-MER database under accessions numbers 3300029977 (YNP-LCB-024), 3300028675 (YNP-LCB-003), 3300005860 (YNP-WB), 3300020139 (GBS), 3300021469 (GB-1845), and 3300021472 (GB-1867). The nine Culexarchaeia MAGs newly recovered in this study are available in NCBI under BioProject ID PRJNA819097. The JZ-Bin-30 MAG is publicly available through the JGI IMG-MER database accession number Ga0181710.

## Supplementary Information

### Supplementary Files

**File Name**: Supplemental Files 1-8 (Excel file)

**Supplementary File 1:** Lists of single marker copy genes, genomes, and genome identifiers used in the phylogenomic analysis shown in Figure 1A and Supplementary Figure 1.

**Supplementary File 2:** Presence-absence patterns for individual genomes, with genes involved in central information-processing machinery.

**Supplementary File 3:** Full list of genes found in Culexarchaeia MAGs that were used to construct Figure 3.

**Supplementary File 4:** Full list of abbreviations used in Figure 3.

**Supplementary File 5:** Full list of IMG or NCBI accessions for the [NiFe] hydrogenase complexes depicted in Figure 4.

**Supplementary File 6:** Presence and absence of methanogenesis marker proteins in Culexarchaeia and Methanomethylicia (Verstraetearchaeota) MAGs.

**Supplementary File 7:** Metadata and geochemical data for YNP sites 003 and 024.

**Supplementary File 8:** Metadata for Culexarchaeia 16S rRNA genes found in NCBI and IMG databases.

**Supplementary Table 1.** Temperature profiles recorded by Alvin’s heat flow probe on December 24^th^, 2016, during dive 4872. This table illustrates the dynamics of the hydrothermal vent field in Guaymas Basin, which makes an exact determination of the temperature at time of sampling hard. To ensure stable placement of the temperature probe, the probe’s disk had to be buried 2-3 cm in the sediment. The sediment sample from which the metagenome was retrieved was taken approximately 40-60 and 30-50 minutes after temperature profiles 1 and 2 were obtained, respectively. To not interfere with the integrity of the sample, the temperatures were taken 1-2 meters away from where the eventual sample was taken. The sample was located approximately half-way between where the two temperature profiles were taken. Geolocation of sample: 27° 00.684 N, 111° 24.266 W. Water depth: 2000 m. The temperatures most closely reflecting the *in situ* temperature range at the time of sampling are highlighted in bold (53.2 and 83.2 ºC).

|  | Heat-flow probe location 1 |  | Heat-flow probe location 2 |  |  |
| --- | --- | --- | --- | --- | --- |
|  | Time 16:33 | **Time 16:36** | Time 16:44 | Time 16:47 | **Time 16:50** |
| Temp. 1 (7-8 cm) | 76.4 °C | **83.2 °C** | 38.6 °C | 50.7 °C | **53.2 °C** |
| Temp. 2 (~17 cm) | 98.3 °C | 101.8 °C | 58.4 °C | 71.1 °C | 73.3 °C |
| Temp. 3 (~27 cm) | 104.2 °C | 105.6 °C | 74.9 °C | 81.8 °C | 82.8 °C |
| Temp. 4 (~37 cm) | 107.4 °C | 108.1 °C | 88.2 °C | 92.1 °C | 93.1 °C |
| Temp. 5 (~47 cm) | 107.2 °C | 107.7 °C | 98.1 °C | 100.1 °C | 101.3 °C |

**Supplementary Table 2.** Proposed naming scheme for the Culexarchaeia MAGs used in this study. Family, genus, and species designations were decided by considering our phylogenomic tree (Figure 1A), pairwise 16S rRNA gene nucleotide identities, and pairwise genome AAI values. *, indicates genome ID used to designate type species.

| Phylum | Class | Order | Family | Genus | Species | Genome ID |
| --- | --- | --- | --- | --- | --- | --- |
| Culexarchaeota<br>(NCBI)<br>/<br>Thermoproteota<br>(GTDB) | Culexarchaeia | Culexarchaeles | Culexarchaeaceae | Culexarchaeum | yellowstonense | YNP-LCB-24-027 * |
|  |  |  |  |  |  | YNP-LCB-3-016 |
|  |  |  |  |  |  | YNP-WB-040 |
|  |  |  |  |  |  | YNP-WB-062 |
|  |  |  |  |  | jinzeense | JZ-Bin-30 * |
|  |  |  |  |  | nevadense | GBS-70-058 * |
|  |  |  | Culexmicrobiaceae | Culexmicrobium | cathedralense | GB-1867-005 * |
|  |  |  |  |  | profundum | GB-1867-035 * |
|  |  |  |  |  | thermophilum | GB-1845-036 * |
|  |  |  |  |  |  | GB-1867-097 |

### Supplementary figures

**Supplementary Figure 1.**
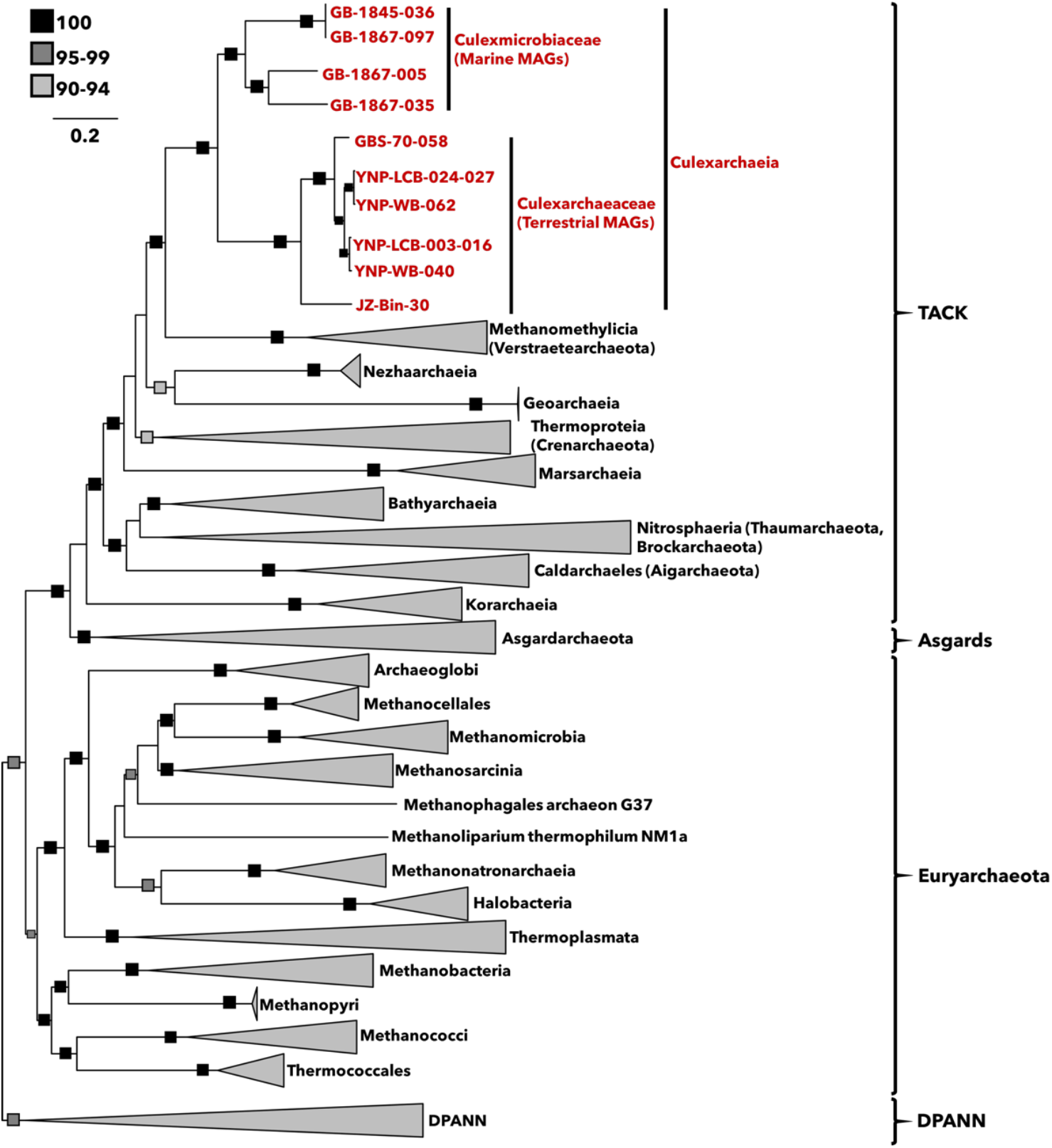
Phylogenomic tree of Culexarchaeia MAGs and reference archaeal genomes. Maximum-likelihood tree, inferred with IQtree and the best-fit LG+C60+F+G model, using a concatenated set of 46 conserved ribosomal proteins (Supplementary File 1). Ultrafast bootstrap support values of 100, 95-99, and 90-94 are indicated with black, dark gray, and light gray squares, respectively.

**Supplementary Figure 2.**
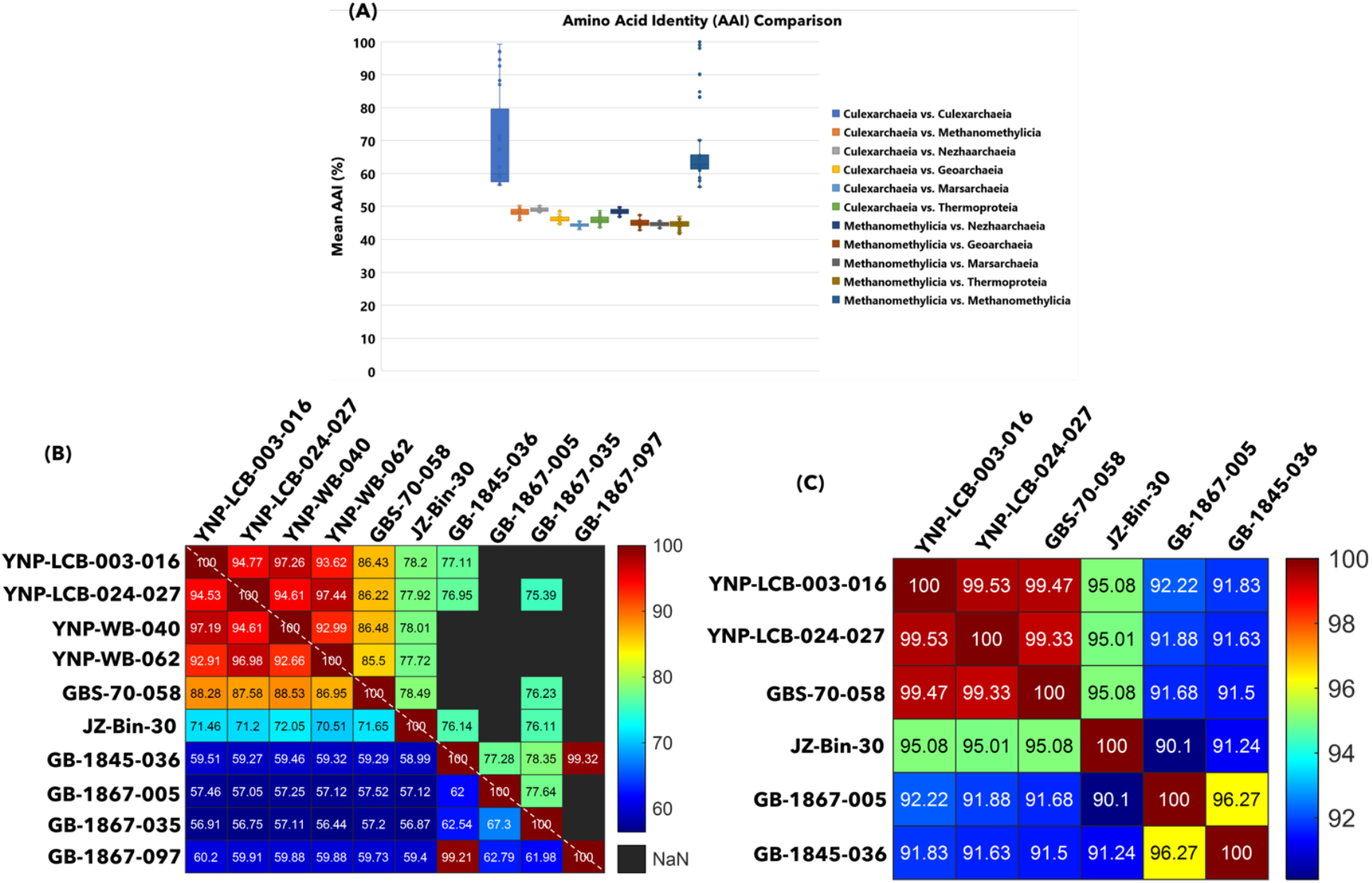
Amino acid identities (AAI) of Culexarchaeia and major TACK lineages. Intra-clade AAI, ANI, and 16S percent identity of Culexarchaeia. **(A)** Analysis shows that the average AAI between TACK classes is below 50%, consistent with the designation of Culexarchaeia as a class level lineage within the TACK superphylum. Thick bar, median; first and third quartile, upper and lower bounds of the box respectively. Upper and lower whiskers extend to the highest and lowest values within 1.5x of the interquartile range, respectively. **(B)** Intra-clade AAI and ANI of Culexarchaeia. Values above and below the dashed white line represent ANI and AAI values, respectively. AAI values were calculated with compareM and ANI values were calculated with fastANI. **(C)** Pairwise % identity of Culexarchaeia 16S rRNA gene sequences calculated by BLASTn.

**Supplementary Figure 3.**
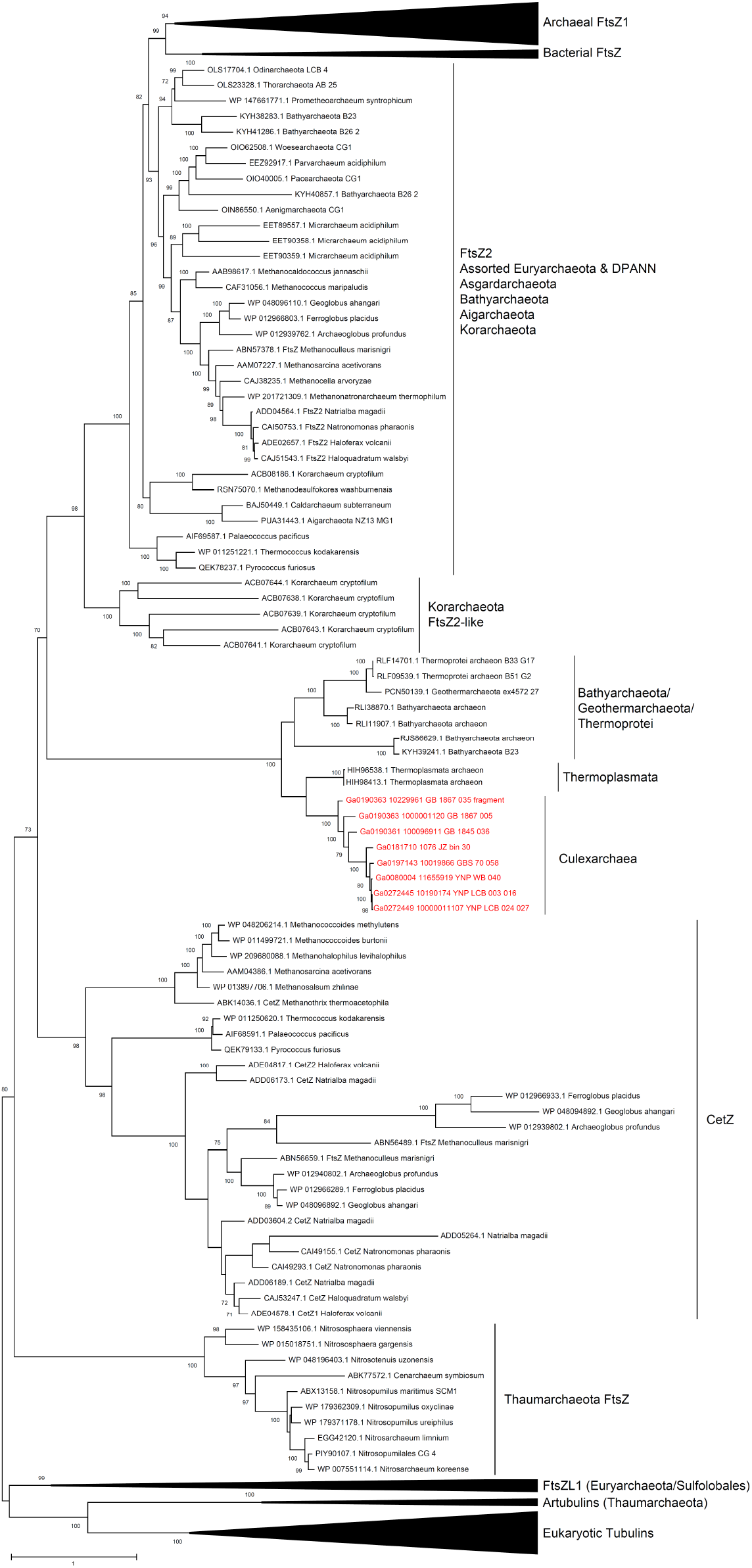
Maximum likelihood phylogeny of tubulin superfamily proteins. Amino acid sequences were aligned using Mafft-LINSi and trimmed with a 70% gap threshold using trimal. Tree was constructed with IQtree2 and the best fit model LG+R5. Ultrafast bootstrap support values above 70 are indicated at each node, and values below 70 are not displayed. Culexarchaeia FtsZ-like sequences are highlighted in red.

**Supplementary Figure 4.**
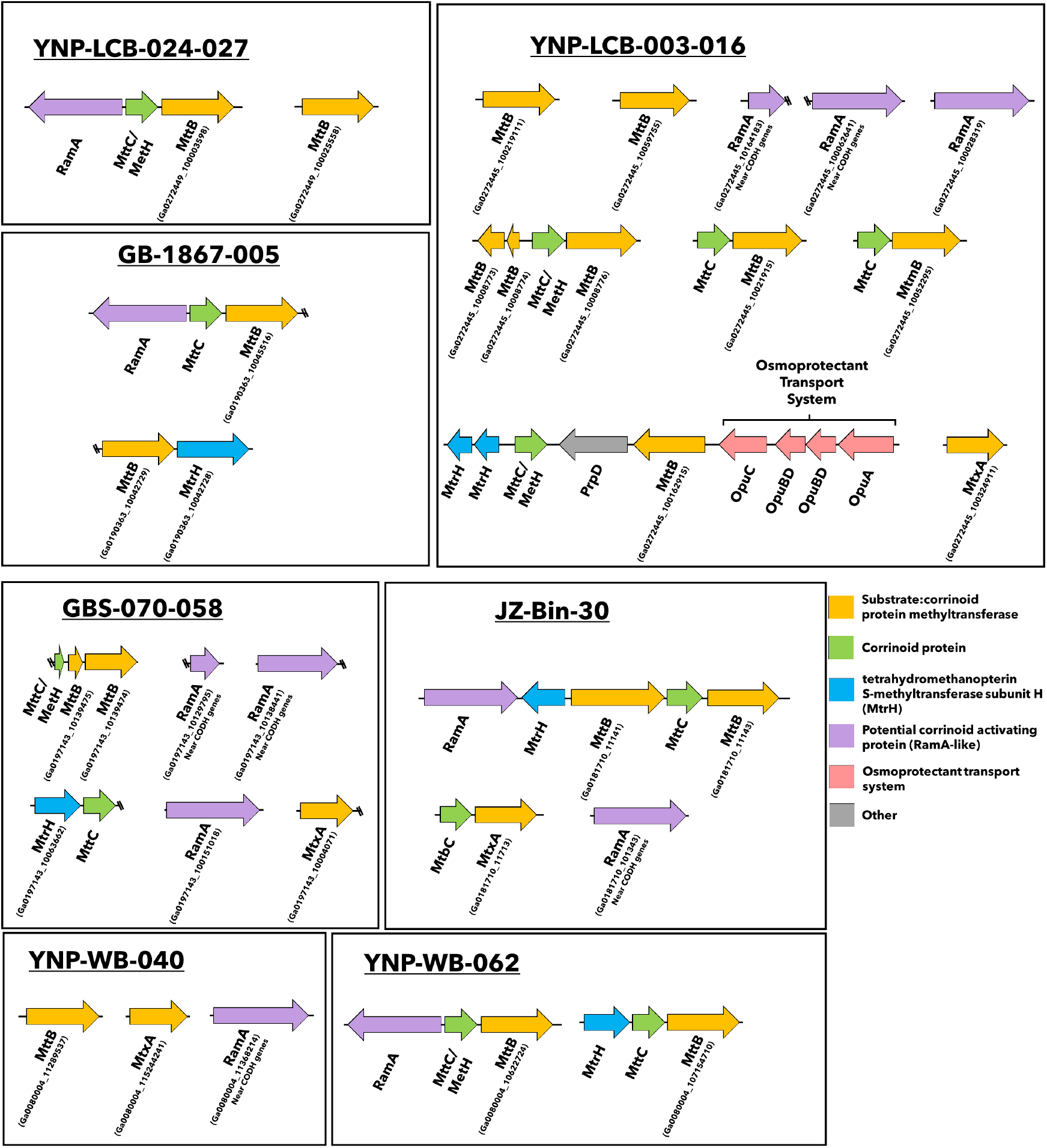
Organization of Culexarchaeia methyltransferase genes. The color of the arrows corresponds to the predicted gene function listed. Some proteins in the family COG3984 (RamA-like) were found to be co-located next to carbon monoxide dehydrogenase (CODH) genes, suggesting they may play a role in the function of this complex instead. Bins not listed did not have any methyltransferase genes encoded. Double slashes indicate the gene is truncated at the end of a scaffold. Each cluster of arrows depicted represents a separate scaffold. Locus tag(s) for genes on IMG are indicated in parentheses for each scaffold.

## Supplementary Methods

### Cell extractions and catalyzed reporter deposition fluorescence *in situ* hybridization (CARD-FISH)

Sediment samples from YNP sites LCB-003 and LCB-024 were collected in July 2019, fixed with paraformaldehyde at a final concentration of 2% for 8 h at 4°C, washed twice with 1x phosphate buffered saline (PBS) and stored in 1x PBS at 4 °C until cells were extracted. Extractions were performed by diluting aliquots of the sediment slurries 1:1 in PBS, adding methanol to a final concentration of 10%, Tween-20 to a final concentration of 0.1%, and vortexing the solution horizontally (Vortex-Genie2, Scientific Industries, Inc., Bohemia, NY) for five minutes to detach cells from particles. After vortexing, an equal volume of 80% (w/v) Nycodenz was layered under the sample and centrifuged at 16,000 g for 30 min at 4°C. Following centrifugation, the supernatant and interphase layers were transferred to a new tube, diluted 1:1 in PBS, and centrifuged at 16,000 g for 5 min to pellet the cells. The supernatant was removed, and the cells were resuspended in 1x PBS and stored at 4°C. Aliquots of the cell suspension were immobilized onto glass slides and CARD-FISH was performed as described in (1) with the following modifications. Cell permeabilization was attempted with either proteinase K (15 µg/mL) for 10 minutes or with 0.1 M HCl for 1 minute at room temperature. Endogenous peroxidases were inactivated by incubating in 0.01 M HCl for 15 minutes. A Culexarchaeaceae-specific probe, Culex824 (5’-TCCACCTAACACCTAGCC-3’), targeted at the 16S rRNA of Culexarchaeaceae found in terrestrial hot springs, was designed using the probe design tool within the Arb software and the Silva 132 reference database (2, 3). Hybridization of probe Culex824 was attempted at formamide concentrations 0, 10, 20, and 35 %. Positive and negative control hybridization reactions were performed with the general archaeal probe Arc915 (5’-GTGCTCCCCCGCCAATTCCT-3’) and NON338 (5’-ACTCCTACGGGAGGCAGC-3’), respectively (4). All probes were labeled with horseradish peroxidase (HRP) and were purchased from Biomers (Ulm, Germany). Tyramide signal amplification was performed with Alexa Fluor 594 (Thermo Fisher). Samples were stained with DAPI, embedded in citifluor, and visualized under an epifluorescence microscope (Leica DM4B).

## Supplementary results and discussion

### Reclassification of JZ-bin-30

Based on our analyses, we propose to reclassify bin JZ-bin-30, originally given the provisional taxonomic assignment *Candidatus* Methanomedium jinzeense (5), as *Candidatus* Culexarchaeum jinzeense sp. nov. (etymology below). We base this re-assignment on the observation in 16S and phylogenomic analyses that JZ-bin-30 does not fall within the candidate class Methanomethylicia, but rather within the new candidate class Culexarchaeia. Additionally, in contrast to what the name Methanomedium implies, all currently available Culexarchaeia genomes, including JZ-bin-30, lack the key enzyme for methanogenesis (methyl-coenzyme M reductase; Mcr) and lack methanogenesis marker genes that have been identified in both *bona fide* Euryarchaeotal methanogens and newly proposed *mcr*-encoding archaea outside the Euryarchaeota (6).

